# The antimycotic 5-fluorocytosine is a virulence inhibitor of uropathogenic *Escherichia coli* and eradicates biofilm-embedded bacteria synergizing with β-lactams

**DOI:** 10.1101/2024.11.20.624304

**Authors:** Srikanth Ravishankar, Antonietta Lucia Conte, Stacy Julisa Carrasco Aliaga, Valerio Baldelli, Karen Leth Nielsen, Maria Pia Conte, Paolo Landini, Elio Rossi

## Abstract

**Objectives:** Biofilm increases bacterial antibiotic tolerance, posing a challenge for treating biofilm-associated infections in clinical settings. 5-fluorocytosine (5-FC), an FDA-approved antifungal and antitumor drug, has been shown to inhibit virulence factor production and biofilm formation in Gram-negative bacteria. This work aims to determine whether 5-FC antivirulence and antibiofilm activity are preserved in clinical *E. coli* isolates and to test possible synergies with antibiotics in treating preformed biofilm.

**Methods:** 5-FC ability to inhibit biofilm formation, disrupt mature biofilm, and modulate virulence determinants was tested on uropathogenic *E. coli* (UPEC) clinical strains using the Crystal Violet-based biofilm adhesion and minimum biofilm eradication concentration assays. Gene expression was measured by RT-qPCR. The effect on bacterial viability within preformed biofilm was monitored using fluorescein diacetate and by determining colony-forming units in biofilm treated with or without 5-FC and specific antibiotics. Bladder epithelial cell cultures were used to assess the effect of 5-FC on UPEC adherence and cytotoxicity.

**Results:** 5-FC inhibited biofilm formation in all tested UPEC strains. Gene expression analysis suggested that 5-FC concurrently inhibits the expression of genes encoding curli fibers, a major adhesion factor in *E. coli*, as well as other virulence determinants such as secreted toxins and type I and P fimbriae. Accordingly, 5-FC reduced UPEC adherence to epithelial cells and promoted host cell survival. Although 5-FC could not disrupt preformed biofilm, combining 5-FC with β-lactams drastically reduced the viability of biofilm-resident bacteria.

**Conclusions:** 5-FC shows antibiofilm and antivirulence activity against uropathogenic *E. coli* strains by reducing the expression of several virulence factors and the overall UPEC pathogenic potential. In combination with β-lactams, 5-FC eradicates bacteria within mature biofilms, which are known to be highly refractory to antibiotic treatment. Our data suggest that 5-FC is an excellent candidate for preventing and treating bacterial infections associated with recalcitrant biofilms.

## INTRODUCTION

Urinary tract infections (UTIs) are a widespread public health concern in community and hospital settings [1]. *Escherichia coli* is the leading pathogen behind UTIs worldwide [2]. Notably, uropathogenic *E. coli* (UPEC) is the most frequent type of *E. coli* that can infect tissues outside the intestines (ExPEC) [3].

The success of UPEC in establishing the infection depends on the production of several virulence determinants. Adhesins, such as curli fibers [4], type-1 fimbriae [5], and P-fimbriae [6], contribute significantly to UPEC ability to cause UTIs. These adhesion factors are essential for forming intracellular bacterial communities (IBCs) within the superficial urinary tract epithelial cells. IBCs allow bacteria to evade the immune system and multiply undetected inside bladder cells, contributing to persistent and recurring infections [7]. In addition, UPECs can secrete soluble toxins that manipulate and damage host cells. The exotoxin α-hemolysin (*hlyA*) promotes lysis of red blood cells and cytotoxic damage to the host uroepithelium [8]. Similarly, secreted toxins termed Serine Protease Autotransporters of *Enterobacteriaceae* (SPATEs) encoded by genes like *sat, pic*, and *vat* have immunomodulatory activity, stimulate vacuole formation, and lead to urothelial barrier dysfunction in bladder and kidneys [9–11].

Antibiotic resistance is another crucial aspect of UPEC infections, particularly associated with recurrent UTIs [12]. Antimicrobial resistance and recalcitrating infection are further exacerbated by the formation of bacterial biofilms [13]. Unlike planktonic bacteria, biofilms are dense, organized, diverse microbial communities [14,15] encased in self-made extracellular polymeric substances (EPS). As biofilms, bacteria exhibit a remarkable resilience against traditional antibiotics [16,17], and their treatment poses significant challenges. For example, transition to a slow-growing or dormant state within biofilms reduces the susceptibility to antibiotics that target cell division [18]. Further, the EPS hinders the penetration of antimicrobial and immune cells [19]. Thus, novel drugs or adjuvants that target biofilm or overcome issues connected with antibiotic resistance are much of a priority.

Recently, we showed that the FDA-approved antimycotic drug 5-fluorocytosine (5-FC) is endowed with antibiofilm activity in the *E. coli* reference strain MG1655 [20]. Here, we demonstrate that the effect of 5-FC is conserved in clinically relevant UPEC isolates obtained from uncomplicated UTIs. Furthermore, we showed that the biological activity of 5-FC is not restricted to biofilm inhibition. The nucleobase analog acts as a potent virulence factors inhibitor in UPEC, repressing the expression of several virulence determinants, hampering hemolysis, and reducing bacterial adhesion and cytotoxicity against bladder epithelial cells. Additionally, we prove that 5-FC synergizes with β-lactams, eradicating bacteria within biofilms.

## MATERIALS AND METHODS

### Bacterial strains and growth conditions

The strains used in this work are reported in Table S1. Nine uropathogenic *E. coli* clinical isolates and one *E. coli* isolate from the fecal flora of a healthy control without UTI from a previously published study were included [21]. These isolates were selected to represent various genotypes of UPEC and were included based on a positive screening for the presence of the curli-encoding operons (*csgBAC* and *csgGDEF)*. For the assays, bacteria were routinely grown in YESCA medium (10 g/L Casamino acids, 1.5 g/L Yeast extract, 0.05 g/L Magnesium sulphate, 0.005 g/L Manganese chloride) at 30°C or 37°C in shaking conditions (150 rpm) or LB agar medium (10 g/L tryptone, 5 g/L yeast extract, 5 g/L sodium chloride, 15 g/L agar). Artificial urine medium (AUM) was prepared according to [22] and adjusted to a pH of 6.4 with NaOH or HCl. To test the effect of nucleobase analog, 5-fluorocytosine (5-FC; stock concentration: 10 mg/ml in sterile milli-Q H_2_O) was added at specified concentrations.

### Gene expression analysis

Gene expression levels were measured using quantitative real-time RT-qPCR as previously described [20]. RNA was extracted from cultures grown in YESCA medium with or without 5-FC at 30°C or 37°C in shaking after 24 hours of incubation. The complete list of primers used for amplification is reported in Table S2.

### Phenotypic assays and bacterial viability determination in mature biofilm

For static biofilm adhesion assays, overnight *E. coli* cultures grown in YESCA medium were normalized to OD_600_ = 0.025 and incubated in the YESCA medium in a 96-well flat bottom plate for 24 hours at 30°C or 37°C in the static condition. Adhesion was determined using the Crystal Violet (CV) staining method, as previously reported [20].

The minimal biofilm eradication concentration (MBEC) of 5-FC and its combination with antibiotics on preformed UPEC biofilm was evaluated. Simultaneously, a fluorescein diacetate (FDA)-based bacterial viability assay and colony-forming units (CFU) determination were performed under the same conditions on mature UPEC biofilms (see Supplementary Methods).

### Hemolysis activity assay

The hemolysis assay was performed on hemolysin-positive UPEC clinical strain KTE223, with hemolysin-negative strains as controls (non-pathogenic *E. coli* strain MG1655 and UPEC clinical strain KTE221). Protocol details are mentioned in Supplementary Methods.

### Infection studies

Infection assays with UPEC clinical strain KTE223 were performed using a human bladder cancer T24 cell line (ATCC HTB-4) (Manassas, USA). Protocol details are reported in Supplementary Methods.

## RESULTS

### 5-fluorocytosine (5-FC) prevents UPEC biofilm formation via pyrimidine starvation and curli fiber inhibition

In the reference strain MG1655, 5-FC inhibits biofilm formation via transcriptional repression of curli fibers [20]. Curli are important fitness factors in UPEC strains [4,23]. Thus, we evaluated whether 5-FC could also demonstrate antibiofilm activity against UPEC, testing biofilm formation and adhesion in 10 clinical isolates (9 UPEC and one fecal isolate) after treatment with 5-FC. Experiments were performed at 30°C, where curli expression is at its maximum [24], and at 37°C, *i*.*e*., the host temperature. 5-FC was able to cause a 1-to 2.3-fold reduction in the biofilm adhesion of three-quarters of the tested UPEC clinical isolates at both 30°C (Fig. 1A) and 37°C (Fig. 1B) at 2.5 µg/ml, a subinhibitory concentration for *E. coli* [20]. The same activity was preserved in artificial urine medium (AUM), which mimics urine composition (Fig. S1).

**Figure 1.**
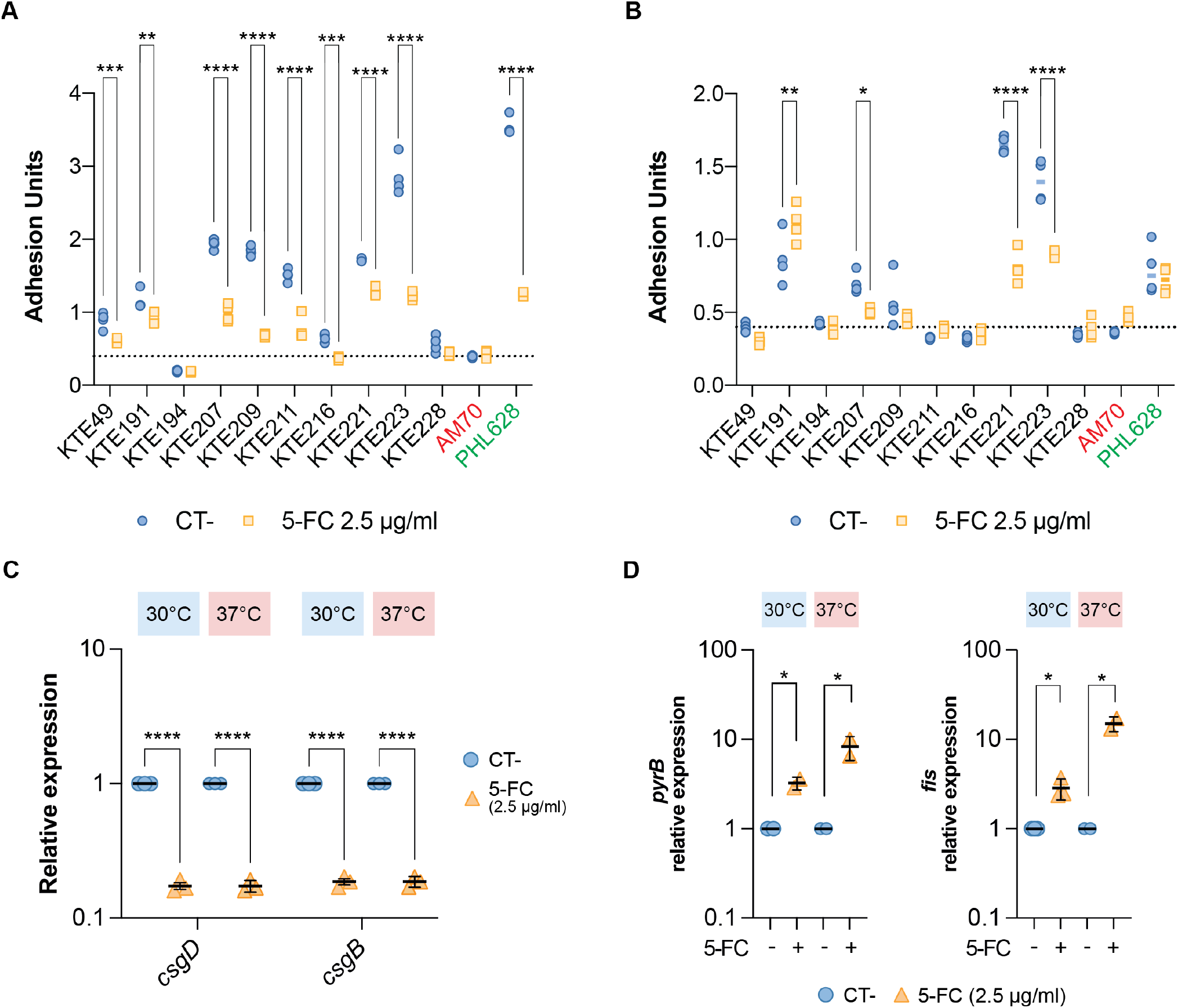
5-fluorocytosine (5-FC) effect on biofilm formation, curli expression, and pyrimidine nucleotide pool in UPEC strains. Bacterial adhesion at **(A)** 30°C and **(B)** 37°C. The assays were performed on 10 UPEC clinical isolates in the presence or absence of 2.5 µg/ml 5-FC. Relative expression of **(C)** curli biosynthetic genes *csgD, csgB*, and **(D)** pyrimidine nucleotide pool-responsive genes *pyrB and fis was* determined by RT-qPCR analysis on RNA extracted from KTE223 UPEC strain in the presence or absence of 2.5 µg/ml 5-FC at 30°C and 37°C. In panels A and B, the dotted line represents the average adhesion of the curli-deficient AM70 strain. In panels C and D, values are expressed as relative units, setting the untreated control to 1. Representative results of 3-4 independent replicates and medians are shown in dot plots. *, p*-*value < 0.05; **, p-value < 0.01; ***, p-value < 0.001; ****, p-value < 0.0001, two-way ANOVA with Dunnett’s test for multiple comparisons.

In the reference strain MG1655, 5-FC-dependent adhesion inhibition is due to the repression of curli-encoding *csgBAC* and *csgDEFG* operons in response to the induced pyrimidine starvation [20]. By focusing on the KTE223 isolate, a strong biofilm producer (Fig. 1A and 1B), we could observe that 5-FC treatment inhibited 5.3 and 5.7 times the expression of *csgB* and *csgD* genes at 30°C and 37°C (Fig. 1C). Further, 5-FC exposure stimulated the expression of *de novo* pyrimidine pathway gene *pyrB* (3.2-fold at 30°C and 8.3-fold at 37°C) and nucleotide-responsive regulator Fis (2.8-fold at 30°C and 15-fold at 37°C) (Fig. 1D), whose expression is activated in response to a decrease in the pyrimidine pool, suggesting that 5-FC treatment leads to pyrimidine starvation in UPEC, similar to what observed in *E. coli* MG1655 [20].

### 5-fluorocytosine (5-FC) inhibits the expression of virulence determinants hindering UPEC pathogenicity

In addition to inhibiting curli gene production, *de novo* pyrimidine biosynthesis perturbation affects type I fimbriae expression in adherent-invasive *Escherichia coli* [25]. Therefore, we reasoned that 5-FC might also negatively impact other virulence factors in UPEC. We tested the expression of type I fimbriae gene *fimA* and several other virulence determinants encoded in the genome of the KTE223 strain, such as P fimbriae (*papC*), α-hemolysin (*hlyA*) and secreted autotransporter toxins (*pic, vat, sat* and *sinH*) upon 5-FC treatment (Fig. 2A and Fig. S2).

**Figure 2.**
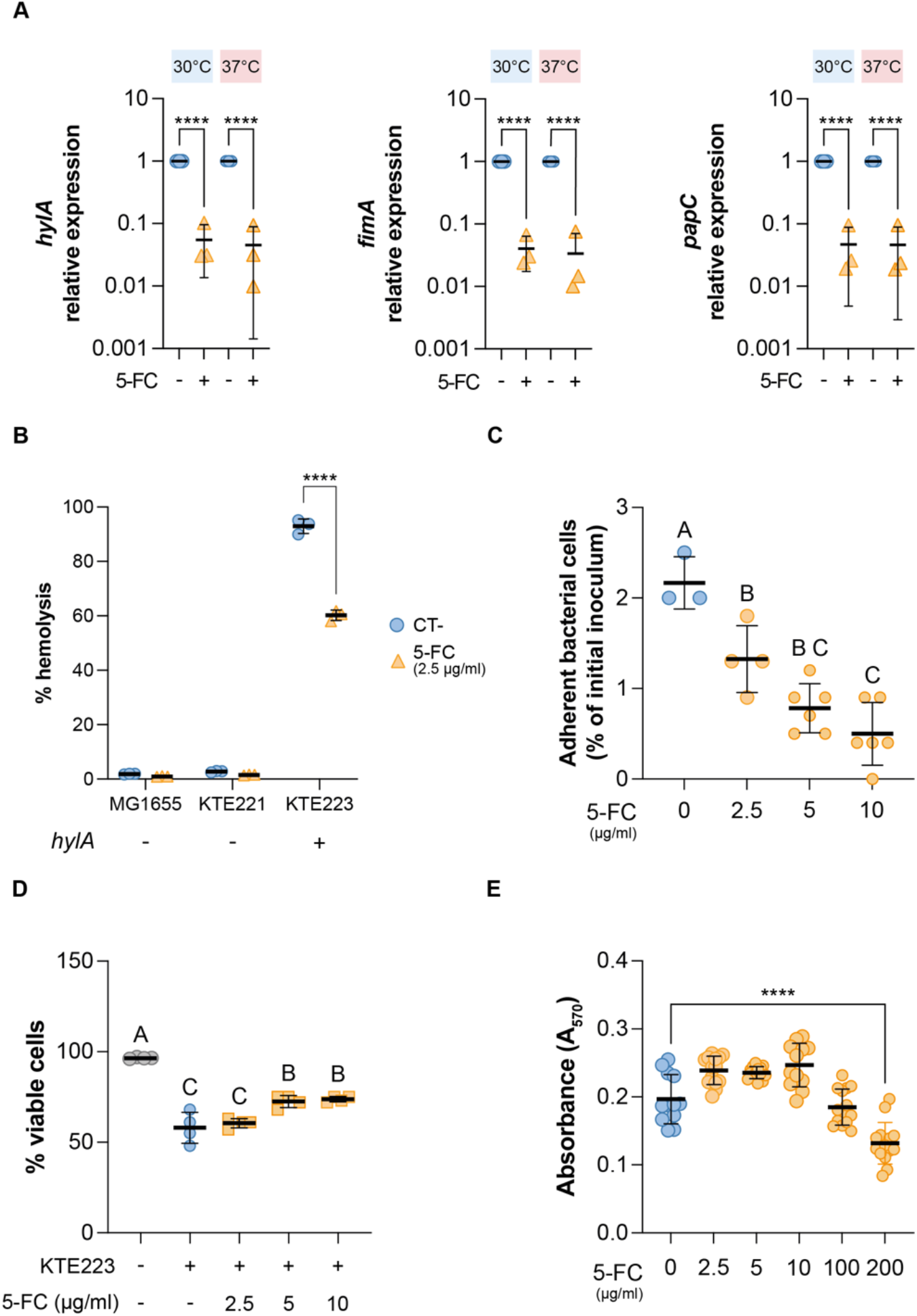
5-FC effect on the expression of virulence factors and pathogenesis of UPEC. **(A)** Relative expression of virulence determinant *fimA, hlyA* and, *papC* determined by RT-qPCR analysis on RNA extracted from KTE223 strain in the presence or absence of 2.5µg/ml 5-FC at 30°C and 37°C. The values are expressed as relative units, setting the untreated control to 1. **(B**) Hemolysis activity of *hlyA*-negative (MG1655 and KTE221) and *hlyA*-positive (KTE223) *E. coli* strains with or without 2.5µg/ml 5-FC. **(C)** KTE223 adhesion to bladder epithelial cells in the presence or absence of increasing concentrations of 5-FC. **(D)** Epithelial cell viability during infection with or without KTE223 and increasing concentrations of 5-FC. Results from the trypan blue exclusion assay are shown. **(E)** Cytotoxic effect of increasing concentrations of 5-FC on bladder epithelial cells after 24 hours of incubation. Results from the colorimetric MTT assay are shown. Representative results of at least 3 independent replicates and medians are shown in all dot plots. *, p*-*value < 0.05; **, p-value < 0.01; ***, p-value < 0.001; ****, p-value < 0.0001 (two-way ANOVA with Dunnett’s test for multiple comparisons). Letters indicate significant within-group differences between treatments (one-way ANOVA with Tukey’s test for multiple comparisons)

5-FC was able to reduce the expression of the adhesin-encoding genes *fimA* and *papC* by 30 and 22 times, respectively, at host temperature (Fig. 2A). Similarly, we observed repression of secreted toxin-encoding genes by 5-FC (Fig. 2A and S2): *hlyA, pic*, and *sinH* expression was reduced 22, 3.6, and 2.6-fold, respectively, at 37°C, while *vat* and *sat* genes showed repression (3.2 and 3.8-fold, respectively) only at 30°C. Consistent with the inhibition of *hlyA* expression, the KTE223 strain showed a 33% reduction in its hemolytic activity when treated with 5-FC at 2.5 µg/ml (Fig. 2B). At the same time, no effect was observed in *hlyA*-negative strains KTE221 and MG1655 (Fig. 2B).

In agreement with our results, treatment with 5-FC of the KTE223 strain reduced bacterial adhesion to epithelial bladder cells by 4.3 times at the highest concentration tested (Fig. 2C). Similarly, 5-FC promoted the viability of epithelial cells in a dose-dependent manner (Fig. 2D), with a maximal effect at concentrations of 5-10 µg/ml, which are sub-inhibitory for bacterial growth [20] and are not toxic to epithelial cells (Fig. 2E) up to 100 µg/ml. Overall, our data indicate that 5-FC activity is not restricted to the inhibition of curli gene expression. Indeed, the molecule can generally affect the production of several virulence determinants important during colonization and acute infection.

### 5-FC potentiates β-lactam antimicrobial activity against bacteria embedded in biofilm

While 5-FC can prevent *E. coli* biofilm formation and adhesion, preformed mature biofilms present a challenge in clinical settings [13]. Thus, using the strong biofilm-producing UPEC strain KTE223, we evaluated the ability of 5-FC to eradicate preformed biofilms and affect bacterial viability within the adherent community. Furthermore, we tested its potential effect when combined with different antibiotics used to treat UTIs [26]. To this end, we measured the Minimum Biofilm Eradication Concentration (MBEC) of 5-FC alone at 2.5 µg/ml, *i*.*e*., the sub-inhibitory concentration able to reduce adhesion (Fig. 1A and [20]), and in combination with increasing concentrations of ciprofloxacin (CIP), gentamicin (GEN), ceftazidime (CAZ), piperacillin (PIP), and aztreonam (ATM). 5-FC treatment and its combination with antibiotics did not disrupt the mature biofilm formed by KTE223 for all the concentrations tested, always showing the same amount of adherent cells as the untreated control (Fig. S3 and Fig. S4). However, when we evaluated bacterial viability within the biofilm using fluorescein diacetate (FDA), we observed that, while the single use of any antibiotics did not result in a reduction of bacterial viability (Fig. 3A and Fig. S5), the combination of 5-FC with β-lactams (CAZ, ATM, PIP) caused a substantial decrease in FDA fluorescence (Fig. 3A). For instance, the combination of 2.5 µg/ml 5-FC with CAZ caused a 50% reduction of fluorescence at 2 µg/ml. This effect was similar at 30°C and 37°C (Fig. 3A), with a slightly higher impact at host temperature. Consistently, the enumeration of living cells within the preformed biofilm after treatment (Fig. 3B) indicated a 50% reduction in CFU/ml upon combining 2.5 µg/ml 5-FC with 2 µg/ml CAZ. Viability reduction was not observed in the presence of 5-FC alone (Fig. S5A) or in combination with fluoroquinolones or aminoglycosides (Fig. S5B). Still, it was preserved in other biofilm-forming isolates (Fig. S6). In addition, bacterial titer in supernatants used for treating mature biofilm showed no differences in CFU/ml in the presence of 5-FC or 5-FC with different CAZ concentrations (Fig. S7), suggesting that biofilm-embedded bacteria are not released into spent media after the drug treatment and that the reduction of viability is due to bacterial cell death.

Overall, our data indicate that 5-FC can specifically synergize with β-lactams, leading to the death of bacteria within biofilms, *i*.*e*., microorganisms known to be refractory to treatment without physically disrupting the biofilm structure.

**Figure 3.**
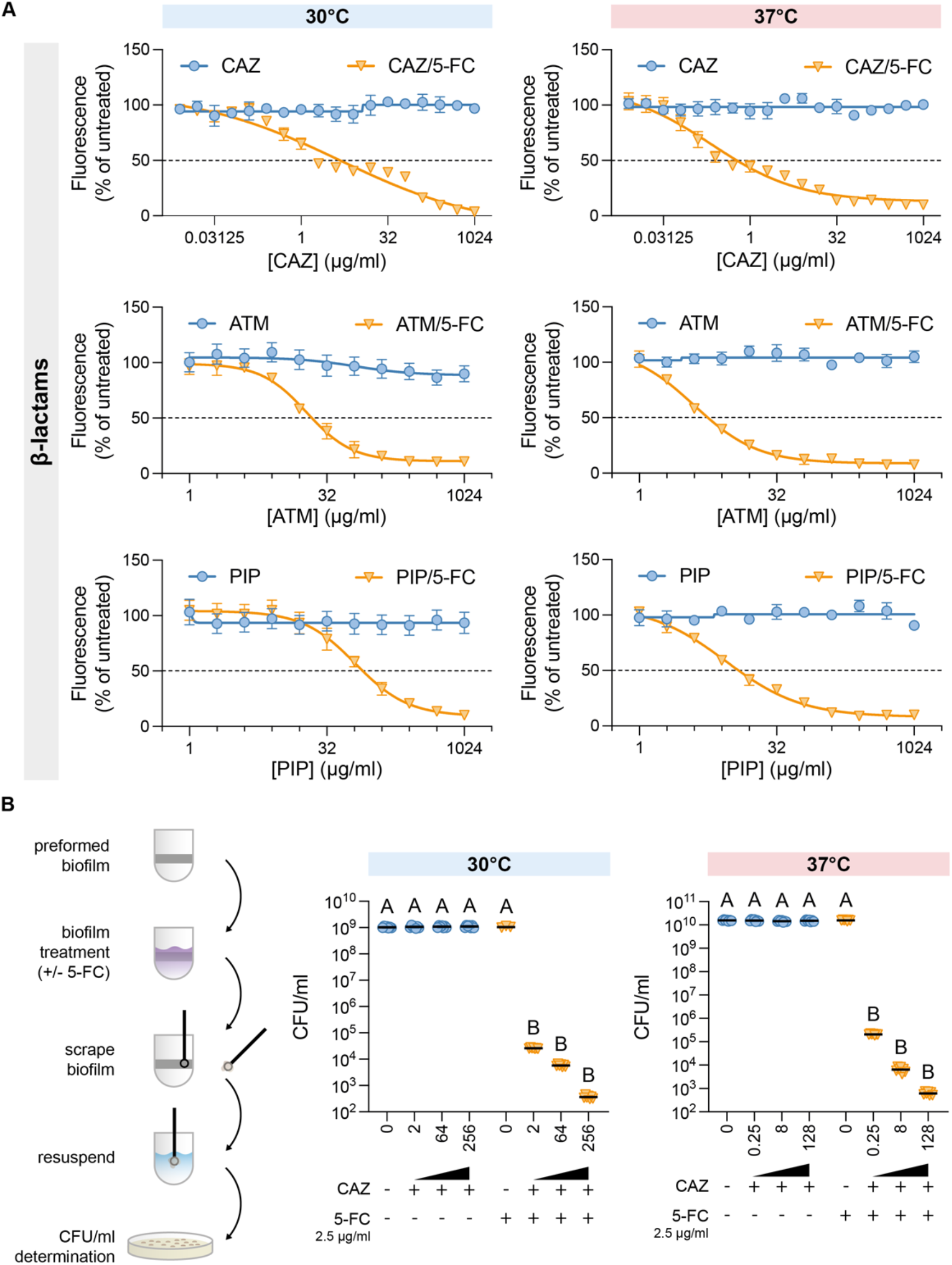
5-FC synergy with β-lactams in mature biofilm of KTE223 strain. **(A)** Metabolically active bacterial cells residing in the mature biofilm of KTE223 strain untreated or treated with increasing concentrations of β-lactams [ceftazidime (CAZ), aztreonam (ATM), and piperacillin (PIP)] and their combination with 2.5µg/ml 5-FC (5-fluorocytosine) at 30°C and 37°C was determined using FDA assay. Fluorescence emitted by the untreated mature biofilm of the KTE223 strain is considered 100%. **(B)** Colony-forming units per ml (CFU/ml) were determined by agar plating method in scrapped biofilm after treating KTE223 mature biofilm with select concentrations of CAZ (ceftazidime) and its combination with 2.5µg/ml 5-FC (5-fluorocytosine) at 30°C and 37°C. The scheme on the left represents the methodology followed to perform this experiment. Results of 3-7 independent replicates and standard deviations are shown. Letters indicate significant within-group differences between treatments (One-way ANOVA with Tukey’s multiple comparisons test).

## Discussion

Bacteria growing in biofilms pose a significant healthcare challenge. Encased within an exopolymeric matrix, they become refractory to antibiotics, rendering current treatments largely ineffective [16,17]. Biofilm-mediated antibiotic tolerance leads to relapsing infections, such as those observed in recurrent urinary infections (UTIs) [13]. Despite extensive research on identifying new antibiofilm drugs [27,28], none have been clinically applied.

In this work, we showed that 5-fluorocytosine (5-FC), an FDA-approved antimycotic drug possessing antibiofilm properties against the *E. coli* reference strain MG1655 [20], exhibits the same activity against diverse clinically relevant *E. coli* strains isolated mainly from UTI. The reduction in biofilm formation is preserved at both environmental and host temperatures (Fig. 1A and Fig. 1B) and in artificial urine medium (Fig. S1), indicating the drug’s applicability in physiologically relevant conditions. Yet, 5-FC’s biological activity is not restricted to biofilm inhibition. The molecule has a broader clinically relevant effect, reducing the expression of several virulence determinants conserved in UPEC. This includes adhesins such as curli fibers, type-1 fimbriae, and P-fimbriae used for colonizing host cells, as well as α-hemolysin and secreted toxins (*pic, vac, sat, sinH*) that induce host tissue damage [8–11]. Repression does not entirely abrogate pathogenicity, but low levels of 5-FC are sufficient to reduce bacterial colonization of the bladder epithelium and promote the survival of host cells (Fig. 2C and Fig. 2D). This suggests that the introduction of 5-FC in clinical practice might be an alternative or, better yet, an adjuvant to antibiotic treatment. Indeed, 5-FC seems capable only of preventing colonization but cannot disrupt preformed biofilm, a condition that often needs to be addressed in clinical practice [13]. Biofilms reduce antibiotic efficacy, and the number of molecules that can directly target bacteria within a biofilm is scarce [29]. In this context, 5-FC shows a peculiar and attractive feature. Indeed, 5-FC was able to potentiate the antibacterial activity of the peptidoglycan biosynthesis-targeting classes of antibiotics (β-lactams/cephalosporins) against *in vitro*-grown biofilm-embedded UPEC (Fig. 3 and Fig. S6). The effect is specific, and 5-FC showed no synergy with ciprofloxacin and gentamicin, which target DNA replication and protein synthesis, respectively (Fig. S5). Although the mechanism of action is currently unknown, 5-FC exhibits antibiofilm activity by affecting intracellular pyrimidine availability in the *E. coli* laboratory strain MG1655, partly via a major peptidoglycan synthase enzyme [20]. Increased expression of genes responding to pyrimidine nucleotide availability upon 5-FC treatment (Fig. 1D) in UPEC and the observation that 5-FC synergizes with drugs targeting peptidoglycan biosynthesis further suggest the existence of a conserved connection between pyrimidine nucleotide biosynthesis and virulence mechanisms in *E. coli* pathotypes, which would not be restricted to UPEC but also apply to other pathogenic strains, as we recently showed [25]. Thus, 5-FC would be helpful in different clinical settings where *E. coli* is the primary pathogen.

Since 5-FC is already used in clinical practice, the concern about bioavailability and safety is low, consistent with our cytotoxicity assay results (Fig. 2E). Thus, repurposing 5-FC could be a transformative approach to address the unmet clinical need for effective treatment to combat biofilm-based infections [30]. It can be combined with β-lactams to eradicate biofilm bacteria or used alone to reduce tissue damage and prevent reinfection by inhibiting virulence factors. Overall, this work has taken us a step forward in effectively introducing 5-FC into clinical practice to improve the efficacy of existing antibiotics against biofilm-associated bacterial infections.

## Transparency declaration

### Contribution

E.R and P.L designed the study with the contribution of M.P.C.; S.R., A.L.C., S.J.C.A. and V.B carried out the experiments; S.R., A.L.C, M.P.C, and E.R. analyzed the data and prepared the figures; K.L.N. provided clinical strains; E.R., P.L., and M.P.C provided financial support; S.R. and E.R. wrote the article; all the authors contributed to the revision of the manuscript.

