## Supplementary Information for "The antimycotic 5-fluorocytosine is a virulence inhibitor of uropathogenic *Escherichia coli* and eradicates biofilm-embedded bacteria synergizing with β-lactams"

**Supplementary Methods**

**Bacterial viability determination in mature biofilm.** The Minimal Biofilm Eradication Concentration (MBEC) of 5-FC and its combination with antibiotics on pre-formed *E. coli* biofilm was evaluated using the standard microdilution method, according to the Clinical and Laboratory Standards Institute guidelines with minor changes [1]. Briefly, overnight cultures were normalized in 200 µL of YESCA to an $OD_{600} \approx 0.025$ in 96-well microtiter plates and then incubated at 30°C or 37°C in static conditions to allow biofilm formation. After 20 hours of growth, the planktonic cells were removed and transferred to a new plate, and the $OD_{600}$ was measured. The pre-formed biofilm was treated with 200 µL of YESCA containing the desired dilutions of 5-FC, antibiotics, or their combinations. The plates were incubated at 30°C or 37°C for 6 hours. The spent media was removed, and biofilm quantification was performed using CV staining method as described earlier. Simultaneously, the same procedure was repeated to determine the viability and metabolic activity of bacteria inside preformed biofilm by measuring the hydrolysis of fluorescein diacetate (FDA) to yellow fluorescent molecule 'fluorescein', according to [2]. Briefly, a solution of 10 mg/ml FDA in acetone was diluted to a final concentration of 0.1 mg/ml in 100mM 3-(N-morpholino)-propanesulfonic acid (MOPS, pH 7.0) and 200 µl of this solution was added to wells with drug-treated or untreated pre-formed biofilm after removing the spent media. After 1 hour of incubation at 37°C in static conditions, the absorbance is read at 494 nm, and the fluorescence is expressed as % of the untreated condition (100%).

For colony-forming units (CFU) determination in preformed biofilm after treatments, biofilm is scrapped using a sterile inoculation loop and resuspended thoroughly in YESCA medium by pipetting vigorously after removing the spent media from the wells. The spent media and the biofilm resuspension are serially diluted and plated on LB agar plates. After overnight incubation at 37°C, colonies were counted, and bacterial titer was expressed as colony forming unit per milliliter (CFU/ml). Untreated pre-formed biofilm was used as a negative control.

**Hemolysis activity assay.** The hemolytic activity was determined as previously described [3,4] with minor modifications. The overnight-grown *E. coli* cultures were centrifuged at 5000 rpm for 10 minutes and resuspended in YESCA medium to reach $OD_{600}$ of 0.2. The cultures ($\sim 10^7$ CFU/ml) were then incubated with 5% (final concentration) defibrinated sheep blood and

10 mM $CaCl_2$ at 37°C in a total volume of 1 mL. 9 g/L NaCl was added to all samples to prevent hypo-osmotic lysis of the erythrocytes. After 2 hours of incubation at 37°C with gentle agitation (300 rpm), intact erythrocytes were harvested by centrifugation (10,000 rpm) at 4°C for 8 minutes. Hemoglobin released in supernatants was determined by measuring the absorbance at 545 nm wavelength (A545). The % hemolysis (P) was calculated using the equation P=[(X-B)/(T-B)]*100, where X is the A545 of the sample analyzed, while B and T represent the baseline and total hemolysis*, i.e.,* the A545 obtained with sterile YESCA (10mM $CaCl_2$ and 9g/L NaCl added) and double-distilled water respectively.

**Cell line cultivation.** The human bladder cancer T24 cell line (ATCC HTB-4) was cultured in Roswell Park Memorial Institute (RPMI) 1640 medium supplemented with 10% heat-inactivated fetal bovine serum (FBS; SAFC Biosciences Inc., Lenexa, KS, USA), 100 IU/L penicillin and 100 mg/L streptomycin. Cultures were maintained in a humidified atmosphere containing 5% $CO_2$ at 37°C.

**Adhesion to T24 cell monolayers.** The adhesiveness of *E. coli* strain KTE223 to T24 monolayers was assayed by culturing the cells in a 24-well plate at a density of $2x10^5$ cells/ml for 24 h at 37 °C in 5% CO2. *E. coli* KTE223 was grown in YESCA with or without 5-FC at 37°C. After overnight growth, bacteria were sub-cultured (1:100) in a fresh medium with or without 5-FC for 2 hours at 37 °C to obtain exponentially grown bacteria.
T24 cells were infected with bacteria in the presence or absence of 5-FC in the cell medium, at a multiplicity of infection (MOI) of approximately one bacterium per cell. They were then centrifuged twice at 500 x g for 2.5 minutes to synchronize infection and incubated for 30 min at 37°C in 5% $CO_2$. After incubation, cells were extensively washed with PBS to remove unattached bacteria, lysed, adding ice-cold 0.1% Triton X-100, and seeded on TSA plates to obtain the total viable count. Bacteria were considered adherent when the mean adhesion index (no. of adherent bacteria/initial inoculum) was ≥ 0.8%.

**Trypan blue exclusion assay**. T24 cell viability, with or without *E. coli* KTE223 and 5-FC, was determined 30 minutes post-infection. Cells were detached by trypsinization, and the number of viable cells in each experimental condition was counted using a trypan blue solution (Sigma-Aldrich, Milan, Italy). One part of 0.4% trypan blue was mixed with one part of the cell suspension. The mixture was incubated for ∼3 min at room temperature. Then, 10 µl of the trypan blue:cell mixture was added to a hemacytometer. The number of unstained (viable) and stained (nonviable) cells was determined. The viability of the control (untreated cells) was regarded as >95%. The percentage of viable cells was calculated as follows:

$$\frac{\text{total number of viable cells per ml of aliquot}}{\text{total number of cells per ml of aliquot}} * 100$$

**Cytotoxicity studies by MTT assay.** T24 cells at $3x10^5$ cells/ml concentration were seeded in 96-well plates and cultured for 24 hours at 37 °C with 5% $CO_2$. Different concentrations of 5-FC (2.5, 5, 10, 100, 200 µg/ml) were added to cell monolayers and incubated for 24 hours. Then, 100 µL of 0.5 mg/ml of 3-(4,5-dimethylthiazol-2-yl)-2,5-diphenyltetrazolium bromide (MTT) reagent was added to each well and plates were incubated at 37°C for 4 hours. Afterward, the dye was eluted with 200 µL of DMSO for 10 min at room temperature and, the optical density at 570 nm was measured using a microplate reader (PerkinElmer, Boston, MA, USA).

**Supplementary Figures**

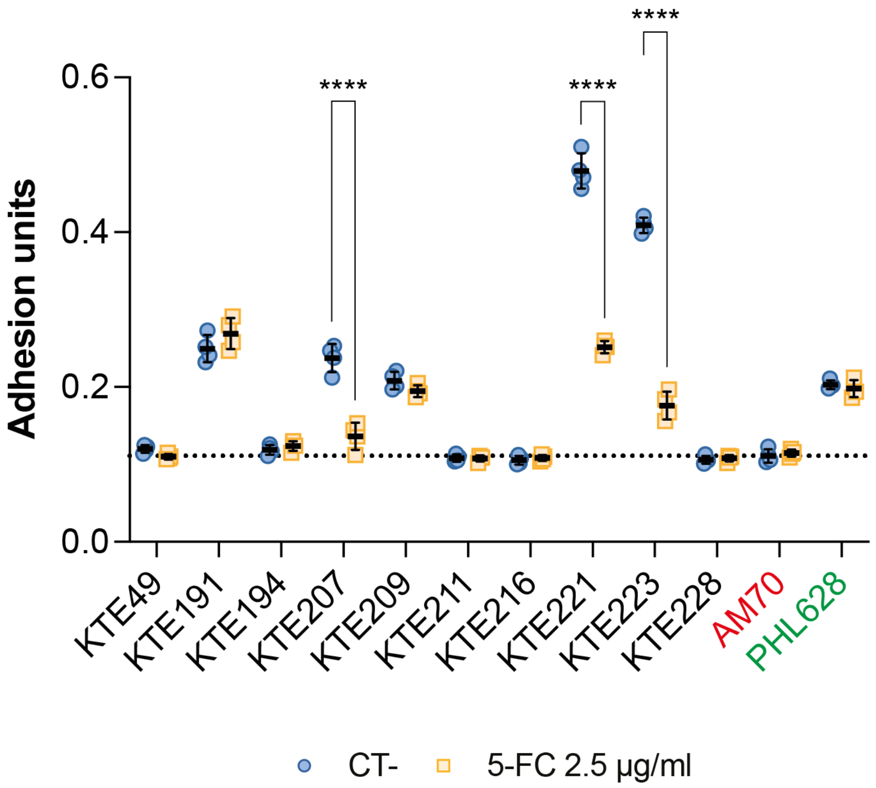

**Figure S1. 5-fluorocytosine (5-FC) effect on biofilm formation against UPEC strains in artificial urine medium (AUM).** Bacterial adhesion in AUM at 37°C is shown. The assay was performed on 10 UPEC strains under study in the presence or absence of 2.5 µg/ml 5-FC. The dotted line represents the average adhesion of the curli-deficient AM70 strain. Representative results of 3-4 independent replicates and medians are shown in dot plots. ****, p-value < 0.0001, two-way ANOVA with Šidák correction for multiple comparisons.

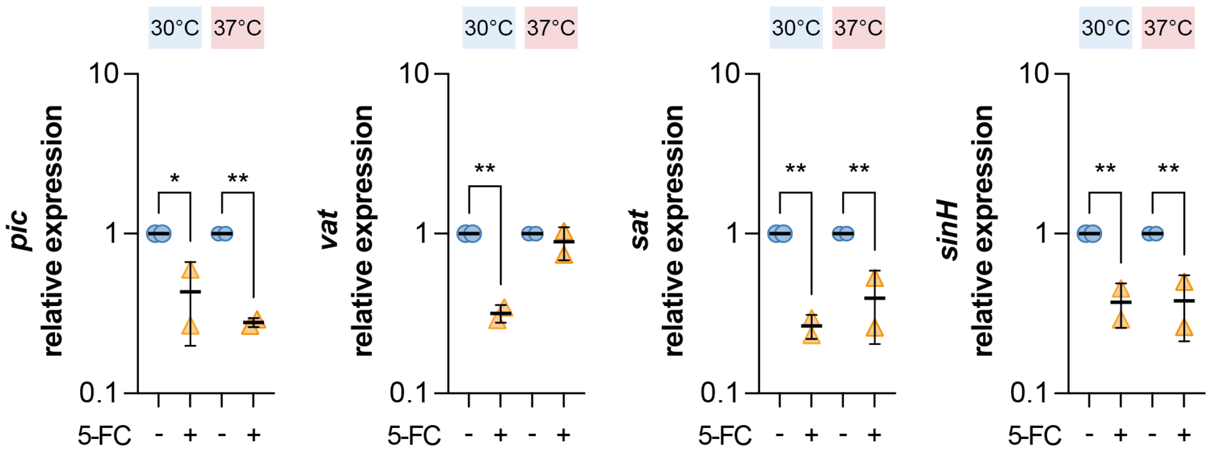

**Figure S2. 5-fluorocytosine (5-FC) effect on transcription of autotransporter toxin genes in KTE223 strain.** Relative expression of autotransporter toxin genes *pic, vat, sat*, and *sinH* determined by RT-qPCR analysis on RNA extracted from KTE223 strain in the presence (triangles) or absence (dots) of 2.5 µg/ml 5-FC at 30°C and 37°C. The values are expressed as relative units, setting the untreated control to 1. The results of 2 independent replicates and medians are shown in the dot plots. *, p-value < 0.05; **, p-value < 0.01, one-way ANOVA with Šidák correction for multiple comparisons.

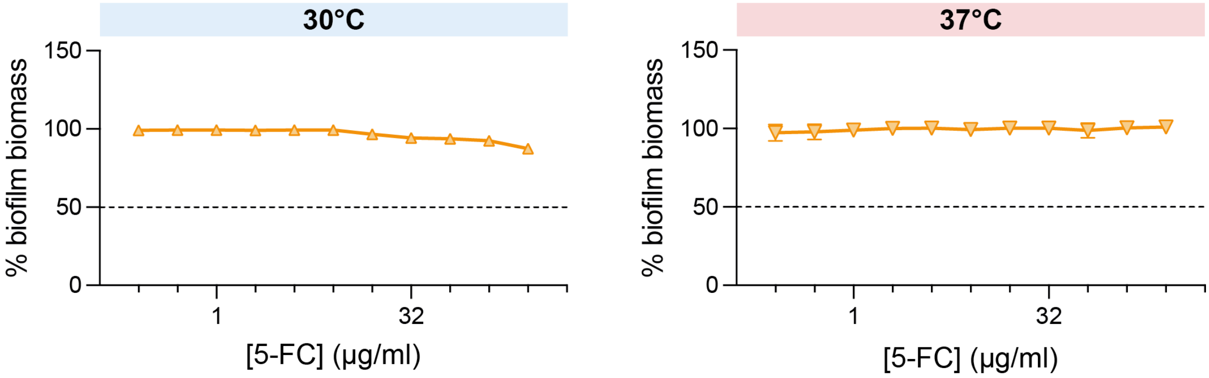

**Figure S3. The effect of 5-fluorocytosine (5-FC) on the mature biofilm of the KTE223 strain.** Percentage of residual preformed biofilm biomass of the KTE223 strain after exposure to increasing concentrations of 5-FC (5-fluorocytosine) at 30°C and 37°C, as determined via crystal violet (CV) staining. CV values in untreated samples were considered 100%. Results from two independent replicates and their standard deviations are shown.

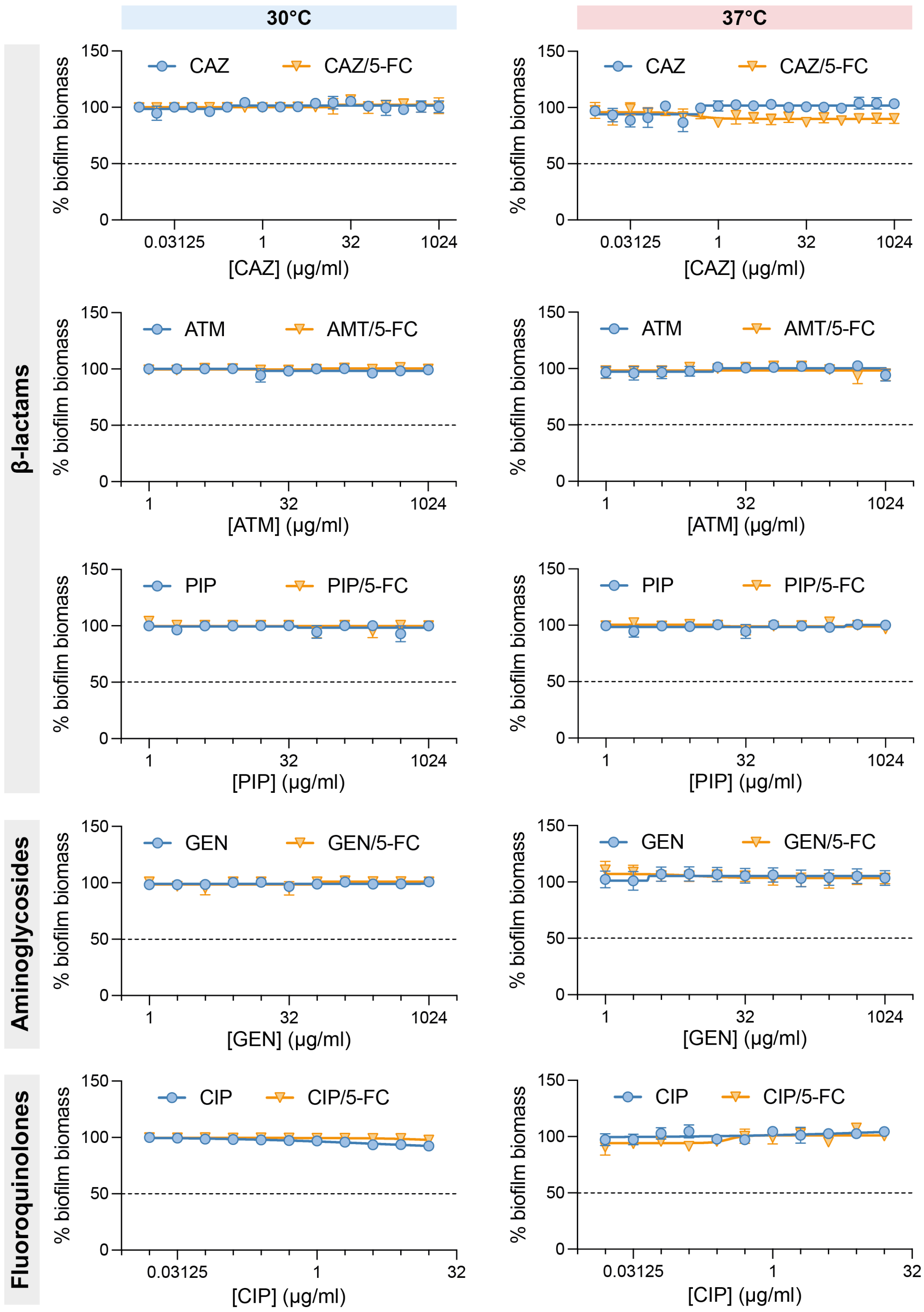

**Figure S4. Effect of antibiotics and their combination with 5-fluorocytosine (5-FC) on mature biofilm of KTE223 strain.** Biofilm biomass of 20-hour preformed biofilm of KTE223 strain untreated or treated with increasing concentrations of various antibiotics and their combination with 2.5 µg/ml 5-FC at 30°C and 37°C was determined. Antibiotics belonging to the class of β-lactams [ceftazidime (CAZ), aztreonam (ATM), and piperacillin (PIP)], aminoglycosides [gentamicin (GEN)], and fluoroquinolones [ciprofloxacin (CIP)] were tested. The untreated preformed biofilm of KTE223 strain is considered 100%. Results of two independent replicates and standard deviations are shown.

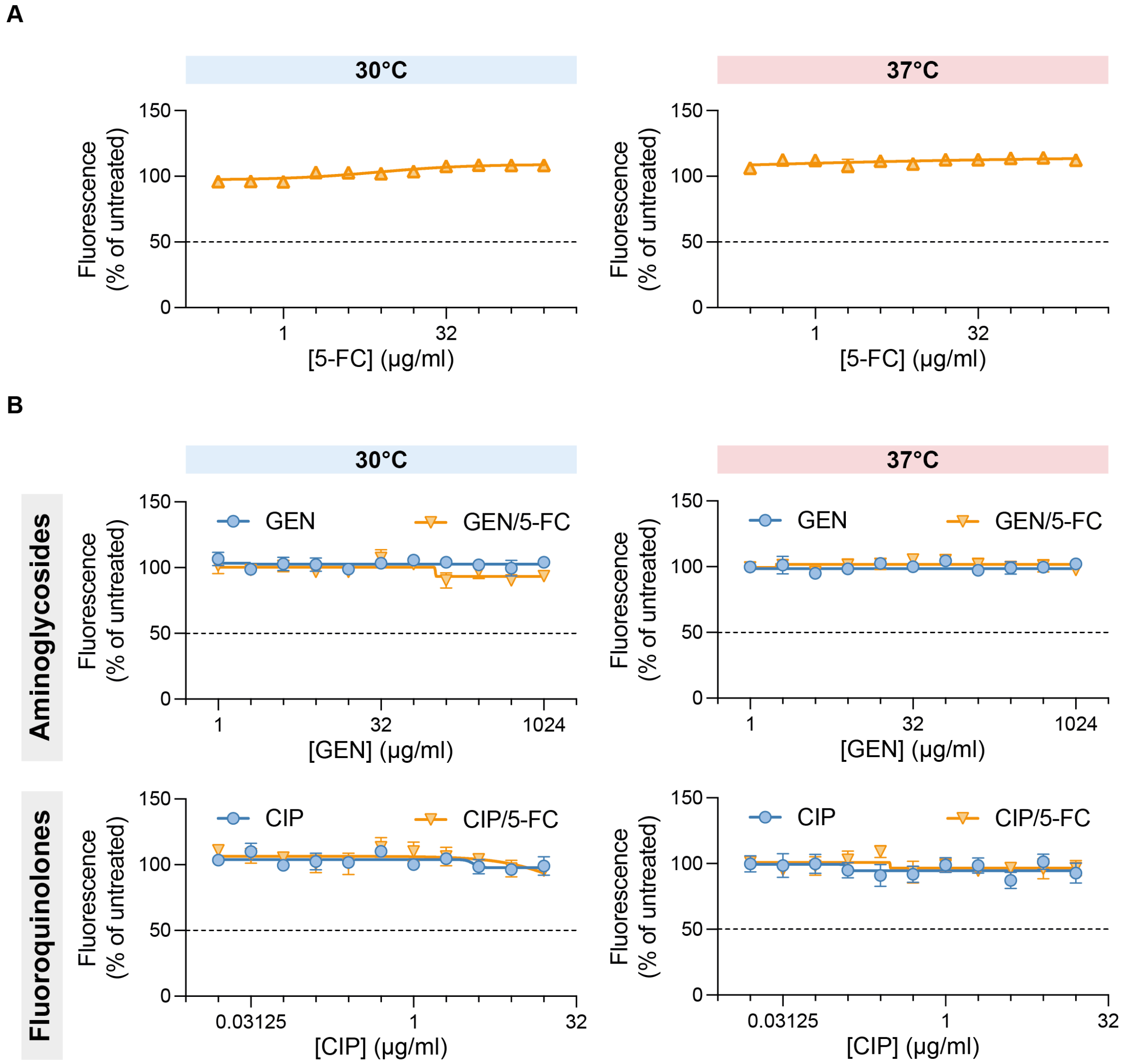

**Figure S5. Effect of 5-FC and its combination with aminoglycoside/fluoroquinolone antibiotics on the viability of biofilm-embedded UPEC bacteria.** Metabolically active bacterial cells residing in the mature biofilm of the KTE223 strain, untreated or treated with increasing concentrations of various antibiotics and in combination with 2.5 µg/ml 5-FC (5-fluorocytosine) at 30°C and 37°C, were determined. Antibiotics belonging to the class of aminoglycosides [gentamicin (GEN)] and fluoroquinolones [ciprofloxacin (CIP)] were tested. The untreated preformed biofilm of the KTE223 strain is considered to be 100%. Results from two independent replicates and standard deviations are shown.

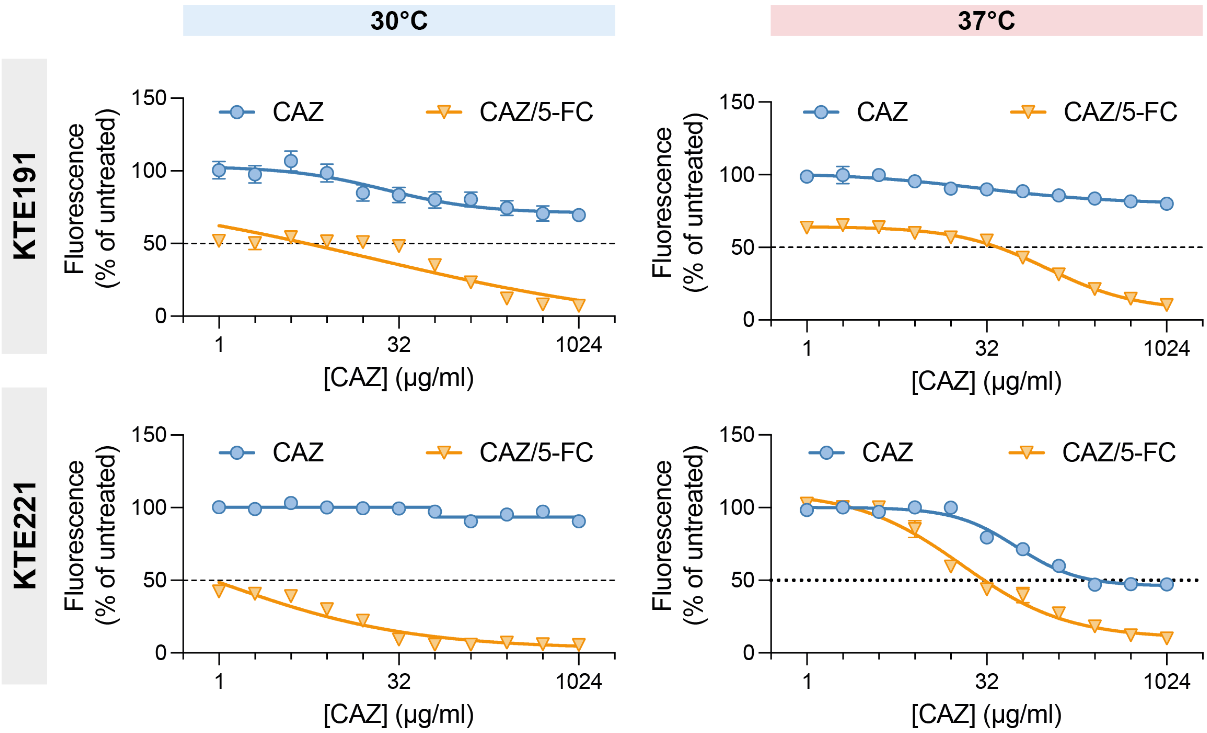

**Figure S6. Effect of 5-FC and its combination with ceftazidime antibiotic on the viability of biofilm-embedded UPEC bacteria.** Metabolically active bacterial cells residing in the mature biofilm of UPEC strains KTE191 and KTE221, untreated or treated with increasing concentrations of ceftazidime (CAZ) and its combination with 2.5 μg/ml 5-FC (5-fluorocytosine) at 30°C and 37°C, were determined. Untreated preformed biofilm of the tested UPEC strain is considered 100%. Results of two independent replicates and standard deviations are shown.

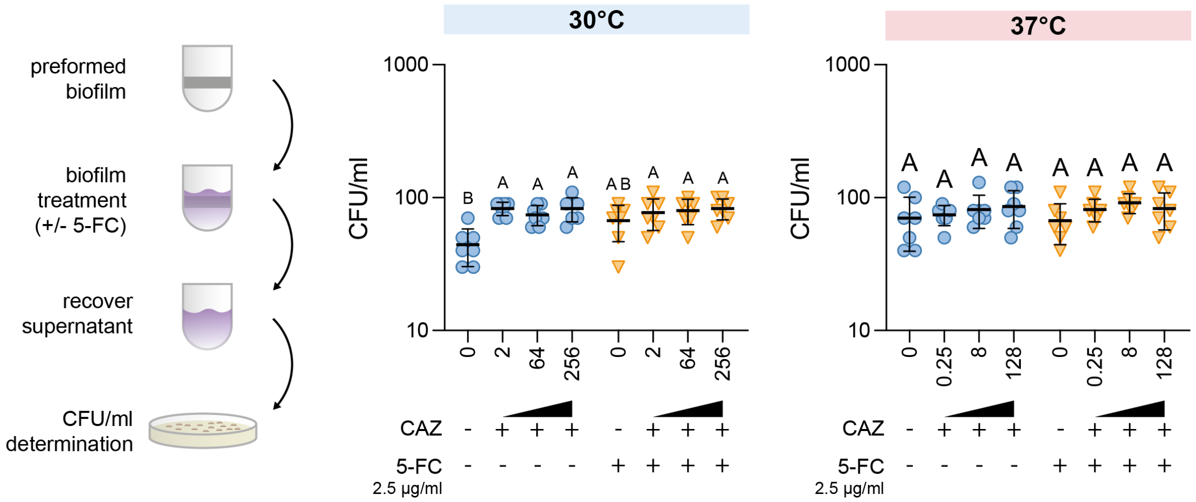

**Figure S7. Viability of bacterial cells in spent media after KTE223 mature biofilm treatment with CAZ and its combination with 5-FC.** Colony-forming units per ml (CFU/ml) were determined by the agar plating method in spent media recovered after treating KTE223 preformed biofilm with select concentrations of CAZ (ceftazidime) and its combination with 2.5 µg/ml 5-FC (5-fluorocytosine) at 30°C and 37°C. The scheme represents the methodology followed to perform this experiment. Results from 7 independent replicates and standard deviations are shown. Letters indicate significant within-group differences between treatments (One-way ANOVA with Tukey's multiple comparisons test).

**Supplementary tables**

**Table S1.** Bacterial strains

| Strain | Relevant genotype | Source | Phylogroup | MLST | Refseq ID | Reference |
|---|---|---|---|---|---|---|
| *E. coli* K-12 str. MG1655 | K-12, F- l- *ilvG*- *rfb*-50 *rph*-1 | Laboratory strain | - | - | - | [5] |
| AM70 (MG1655Δ*csgA*) | MG1655 derivative. Replacement of the *csgA* gene with a chloramphenicol resistance cassette. | Laboratory strain | - | - | - | [6] |
| PHL628 | MG1655 *malA*-Kan *ompR234* | Laboratory strain | - | - | - | [7] |
| KTE49 | Clinical isolate | Fecal | B2 | 131 | GCA_000351445.1 | [8] |
| KTE191 | Clinical isolate | UTI | B2 | 12* | GCA_000351005.1 | [8] |
| KTE194 | Clinical isolate | UTI | B2 | 141 | GCA_000352805.1 | [8] |
| KTE207 | Clinical isolate | UTI | B2 | 998 | GCA_000353005.1 | [8] |
| KTE209 | Clinical isolate | UTI | B2 | 73 | GCA_000353025.1 | [8] |
| KTE211 | Clinical isolate | UTI | B2 | 131 | GCA_000353045.1 | [8] |
| KTE216 | Clinical isolate | UTI | B2 | 131 | GCA_000351225.1 | [8] |
| KTE221 | Clinical isolate | UTI | A | 410 | GCA_000408225.1 | [8] |
| KTE223 | Clinical isolate | UTI | B2 | 73 | GCA_000353125.1 | [8] |
| KTE228 | Clinical isolate | UTI | D | 69 | GCA_000351285.1 | [8] |

*single locus variant

**Table S2.** Primers used for gene expression analysis.

| Name | Sequence (5' - 3') |
|---|---|
| 16S rRNA_RT_frd | TGTCGTCAGCTCGTGTCGTGA |
| 16S rRNA_RT_rev | ATCCCCACCTTCCTCCGGT |
| csgB_RT_frd | CATAATTGGTCAAGCTGGGACTAA |
| csgB_RT_rev | GCAACAACCGCCAAAAGTTT |
| csgD_RT_frd | CCCGTACCGCGACATTG |
| csgD_RT_rev | AAGGAGGGCTGATTCCGTGCTG |
| fimA_RT_for | CTCTGGCAATCGTTGTTCTGTC |
| fimA_RT_rev | TCAACAGAGCCTGCATCAACTG |
| hlyA_RT_fw | ACTCTCATTCCTGTCCATTGCCGA |
| hlyA_RT_rev | GAAGCCAGAACAGTGCTTATCGTTGTTA |
| papC_RT_fw | TACAGTGGCAGTATGAGTAATGACCG |
| papC_RT_rev | GCGGACTACGATGACTGTAATAGGC |
| pic_RT_fw | TGTCCGTTCCGATATTGCCTATCAG |
| pic_RT_rev | GGCCATTGGGGCTTTATCCAGT |
| sat_RT_fw | TCAATTCCGGATTTTTCTGGTGCAG |
| sat_RT_rev | AGAAGACTGAGCGTAAACCTGGG |
| vat_RT_fw | ATCAACGGTTGGTGGCAACAATCC |
| vat_RT_rev | CCATGGGCGCTTTATCAAGATGTCC |
| sinH_RT_fw | GACACACTCTCTCCCTACGGTAAGG |
| sinH_RT_rev | CGTTGCGCAGAAAATTGGCTGAAA |
| pyrB_RT_frd | CGACAGCGCCAATACATCACT |
| pyrB_RT_rev | CGGCATTCAGTACCGGTACAT |
| carA_RT_frd | CGAGAAAGGCGCACAGAATGGC |
| carA_RT_rev | GGCTTCTGCGGTGGTCACTTCT |
| fis_RT_frd | AAAACCCCTGCGTGACTCGGTT |
| fis_RT_rev | CACCATGTCCAACAGGGGCTGT |
